# Effects of Cognitive Demand Reduction on Choice Overload

**DOI:** 10.64898/2026.02.19.706731

**Authors:** Seungbeom Seo, Seungji Lee, Nayoung Lee, Sung-Phil Kim

## Abstract

Choice overload occurs when an ever-growing number of options impairs decision quality, because evaluating options taxes cognitive resources. We investigated whether reducing cognitive demand could mitigate overload by encouraging greater cognitive effort to achieve optimal choice. We conducted two experiments manipulating cognitive demand in complementary ways: Experiment 1 reduced demand by presenting high-attractiveness sets, and Experiment 2 did so by providing a shortlist tool. In both experiments, participants chose from sets of 6–24 options while their eye-gaze and electroencephalographic (EEG) data were recorded. We found that reducing demand made decisions faster, but did not improve choice performance as set-size increased. Under low-demand conditions, eye-gaze measures revealed narrower search and EEG measures showed reduced working memory engagement per option, together indicating less searching and processing efforts. These results suggest that even with reduced cognitive demand, people coast through easier decisions, conserving effort and leaving the choice overload effect largely intact.

## MAIN

Every day, people face a staggering number of decisions – one estimate suggests tens of thousands daily^1^. With such an abundance of decisions, it may seem that having more options is better. Indeed, larger assortments can increase the sense of freedom and variety, potentially making decision-making more enjoyable^2–5^. However, beyond a certain point, an overabundance of options can lead to choice overload, wherein decision-making suffers, producing negative outcomes like stress, regret, or lower satisfaction^6–11^. For example, in digital media, an overabundance of content can cause people to spend more time choosing than consuming (e.g., a “Netflix syndrome”), leading to frustration^12^. In such cases, the cognitive costs of processing many alternatives outweigh the benefits of added variety^13–18^.

A useful framework for understanding choice overload is the evaluation-cost model^13^. As the number of options grows, the mental effort required to evaluate them (cognitive demand) rises sharply, while the marginal benefit of each additional option diminishes. This trade-off suggests that decision-makers would continue examining options only until the expected benefit of doing so no longer justifies the cognitive effort. Neuroscientific evidence supports this notion by showing that reward-related brain regions responded most strongly to moderate assortment sizes, but showed reduced activation for larger choice sets^19^. Thus, when faced with abundant options, people often downshift their cognitive effort to avoid excessive mental cost^20–22^. This tendency to conserve effort can lead to suboptimal decisions, as individuals may not fully explore all options^23,24^.

Since choice overload stems from an overly taxing evaluation process, a logical remedy is to ease the cognitive burden. Because human attentional and working memory capacity is limited, evaluating a large number of options can quickly overwhelm our cognitive resources^25,26^. By reducing the decision-maker’s cognitive demand, people can handle more alternatives without feeling overwhelmed.

In modern consumer environments, two common strategies aim to make choices easier to reduce cognitive demand:

Strategy 1: Preference-Based Filtering (Personalized Recommendations). This approach presents the decision-maker with a curated subset of highly attractive options, tailored to individual preferences. By filtering out less desirable choices (e.g., recommending a short list of top-rated movies), the decision becomes easier because all the options are likely to be good, and the stakes of a “wrong” choice are lower. Thus, the cognitive effort needed to compare options is greatly reduced. For instance, Chernev and Hamilton (2009) found that when consumers were offered only uniformly high-quality options, they perceived less risk and were satisfied with the choice more quickly^27^.

Strategy 2: Sequential Narrowing of Choices. This approach breaks the decision into stages, rather than confronting the chooser with all options at once. A common example is a “wishlist” or bookmarking tool on e-commerce platforms: from a large initial pool, users first mark a subset of favorites, and then make a final selection from that subset. By narrowing the choice set sequentially, the cognitive demand is reduced because each stage involves evaluating fewer options. Importantly, by the final stage, the decision-maker has already evaluated the options during the filtering step, so they are not processing all new information simultaneously. This sequential filtering process allows focus on fewer options at a time, easing cognitive load during the final choice.

However, it remains elusive whether making decisions easier through these industry-driven strategies indeed leads to better choices and prevents overload. On one hand, reducing cognitive demand could enable people to engage more fully with options, avoiding the negative effects of overload. With only high-quality, pre-selected options, individuals might invest their extra mental resources into comparing options more thoroughly, yielding more optimal decisions. On the other hand, simplifying a choice may not guarantee better outcomes. People often act as cognitive misers^28^, seeking to minimize unnecessary mental work. Therefore, when a task is made easier, decision-makers might respond by exerting even less effort, reasoning that minimal effort is now sufficient. Ironically, strategies designed to alleviate overload could cause people to coast through the decision. If so, any benefit of reduced demand is canceled out by reduced effort investment, leaving the choice overload problem intact or even exacerbated.

Therefore, a key research question is whether making a choice set “easier” prompts individuals to reallocate their spare mental capacity to the task or leads them to conserve effort and drift into a low-effort decision strategy. It is difficult to answer this with traditional self-reports or behavioral measures, which are limited in capturing temporal dynamics in the decision process^29^. Instead, leveraging neurophysiological indicators of cognitive effort can reveal how people adjust their engagement when a decision is simplified^30^. Complementing this approach, prior neurophysiological studies of choice overload suggest that excessive alternatives can disrupt early-stage processing and recruit later compensatory engagement^31,32^.

A recent study by Huang and Xu (2025) gave a hint to this question that making choices easier speeds decisions but does not necessarily improve choice performance^33^. Although they did not answer this question by manipulating cognitive demand via task constraints or decision aids, they found that high-preference choice sets significantly accelerated decision processes but also lowered choice performance. In addition, more neural engagement was observed for a small set of options with low-preference (i.e., high-demand), reflecting more mental effort. Yet, increased neural engagement in low-preference small sets did not necessarily yield higher choice performance, warranting clearer explanations of relationships between cognitive demand and effort. Also, set size was either 4 or 16, limiting the understanding of the effect of set sizes, which was fundamental to choice overload problems.

In the current study, we tackle the above question by experimentally testing the two demand-reduction strategies in controlled decision-making tasks. Critically, our study extends beyond preference-based ease by testing a process intervention, sequential filtering via a shortlist tool, and investigates relationships between cognitive demand and effort more deeply by examining both search and processing efforts via a multi-modal approach. Two experiments implementing each strategy were designed to observe how lowering external cognitive demand impacts decision-makers’ effort and choices.

In Experiment 1, we simulated preference-based filtering by manipulating the attractiveness of the choice set, a strategy akin to providing preference-articulated choice set^34^. Participants chose from sets of options that were either all high-preferred (low-demand condition) or all low-preferred (high-demand condition). This mirrors a personalized recommendation system that boosts option attractiveness to ease decision-making. In Experiment 2, we implemented sequential narrowing. Participants first completed a preliminary bookmarking step to select a handful of favorites from a larger set, and then made a final choice from those bookmarked items. In the low-demand condition, the final choice set was “screened” by their bookmarks, whereas in the high-demand condition, the final set was a control assortment without such pre-selection. Detailed task sequences and demand manipulations are illustrated in Supplementary Fig. 1. We also confirmed manipulation checks of the demand conditions (Supplementary Fig. 2).

Notably, we varied the number of options in both experiments (sets of 6, 12, 16, or 24) to examine the interaction of demand reduction with set size. This variety of set sizes allowed us to observe conventional choice overload patterns and see if the interventions alter those patterns^10,19,35^. If reducing cognitive demand encourages greater engagement, we expected less decline in decision quality as the set size increases in the low-demand than the high-demand conditions. Conversely, if participants in easier tasks tend to slack off, there would be no improvement in choice outcomes despite the simplification.

To address these questions, we measured not just which options participants chose, but how they made their decisions. We used multi-modal measurements, including eye-tracking to index visual search behavior and electroencephalography (EEG) to gauge neural engagement, on a “moment-to-moment” basis. For visual search, we extracted three eye-gaze measures: screening ratio, revisit ratio, and dwell bias^36–39^. Screening ratio indexes the exhaustiveness of initial search – the proportion of available options that were fixated at least once. Revisit ratio indexes comparative deliberation – the tendency to return to previously viewed options for additional comparisons. Dwell bias indexes the distribution of attention across items – the extent to which gaze time is concentrated on a subset of options. Together, these metrics capture the search effort exerted by participants to explore the assortment.

For neural processing, we analyzed Fixation-Related Potentials (FRPs), focusing on peak latencies of early posterior N1 (attentional orienting)^40^, occipital P2 (perceptual evaluation/decision processing)^41^, and frontal N2 (conflict monitoring/cognitive control)^42,43^. We also analyzed Fixation-Related Spectral Perturbations (FRSPs) in the occipital alpha band (8-13 Hz), where stronger alpha suppression (i.e., desynchronization) indicates greater attentional engagement^44–48^. Longer FRP latencies and stronger alpha desynchronization are taken to reflect more effortful processing for each fixated option^49^. These fixation-related neural measures provided insight into whether or not participants in the low-demand conditions were truly examining individual options more extensively. In summary, our use of the demand-reduction strategies and fine-grained effort measures tests whether making choices easier leads to effort reallocation or effort conservation.

The primary behavioral outcomes were defined at the trial level and summarized within each Demand × Set Size condition. Choice performance quantified the extent to which the selected option approached the best available alternative in the set, based on each participant’s own pre-task ratings; for each trial, we computed the difference between the ratings of the chosen item and the highest-rated item among the displayed options. Decision time was defined as the latency from choice-set onset to the mouse click indicating selection. These trial-wise measures were then averaged across trials within each condition for each participant (see Methods for full operational definitions and preprocessing details).

### Experiment 1 (Preference-based Filtering)

In Experiment 1, choice performance was significantly influenced by both demand condition and set size. Overall, performance was lower in the low-demand than in the high-demand conditions (*F*_1,16_ = 13.66, *P* < 0.01, η^2^p = 0.46). We also observed an overall decline in performance as set size increased (*F*_3,48_ = 11.07, *P* < 0.001, η^2^p = 0.41). Importantly, a significant Demand × Set Size interaction effect (*F*_3,48_ = 11.89, *P* < 0.001, η^2^p = 0.43) indicated that performance in the low-demand condition worsened substantially as set size increased, while performance in the high-demand condition remained largely stable (Fig. 1a, left). Thus, making options more appealing did not enable participants to maintain high-quality choices as the assortment grew.

**Figure 1.**
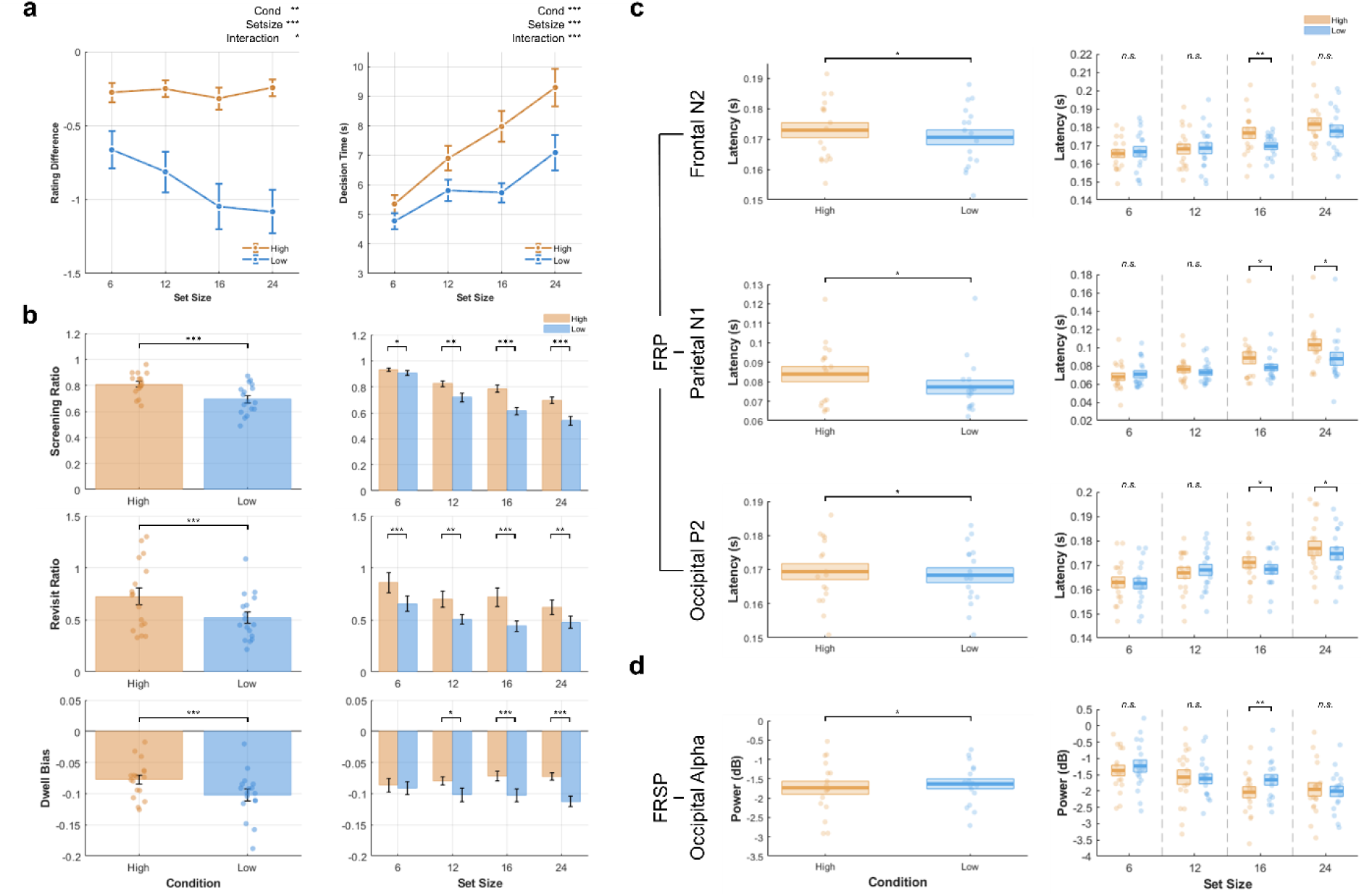
Results of Experiment 1 (Preference-based filtering). **a**, Rating difference between the chosen item and the best-rated item in the set (left) and decision time (right), shown by demand condition (high vs. low) and set size (6, 12, 16, and 24). **b**, Eye-tracking measures – screening ratio (fraction of items viewed), revisit ratio (proportion of items fixated more than once), and dwell bias (normalized gaze-time bias) – shown by demand condition (left; individual participants, n = 17) and by set size (right). **c**, EEG FRP peak latencies **d**, EEG FRSP in the alpha band. In panels c-d, individual participants are shown (n = 17), and box plots summarize the mean (center line) with boundaries indicating ±SEM. Statistical significance from two-way repeated-measures ANOVA is indicated as: *n.s.* for non-significant, * for *P* < 0.05, ** for *P* < 0.01, and *** for *P* < 0.001.

Next, decisions were made significantly faster in the low-demand than in the high-demand conditions (mean 5.85 ± 1.84 s vs. 7.38 ± 2.44 s; *F*_1,16_ = 28.28, *P* < 0.001, η^2^p = 0.64). As expected, larger choice sets took longer overall to decide upon (*F*_3,48_ = 61.33, *P* < 0.001, η^2^p = 0.79). There was also a significant Demand × Set Size interaction (*F*_3,48_ = 10.59, *P* < 0.001, η^2^p = 0.40), reflecting that decision time increased more steeply with set size in the high-demand condition (Fig. 1a, right). In essence, making the choice set easier with high-attractiveness options sped up decisions, but at the cost of accuracy.

Eye-gaze analysis confirmed that participants invested less visual search effort in the low-demand condition. Across all three gaze metrics, values were significantly lower with low demand than with high demand (Screening ratio: *F*_1,16_ = 30.22, *P* < 0.001, η^2^p = 0.65, Revisit ratio: *F*_1,16_ = 24.55, *P* < 0.001, η^2^p = 0.61, Dwell bias: *F*_1,16_ = 18.61, *P* < 0.001, η^2^p = 0.54) (Fig. 1b, left). As set size increased, the screening ratio dropped markedly (*F*_3,48_ = 190.05, *P* < 0.001, η^2^p = 0.92), and so did the revisit ratio (*F*_3,48_ = 16.47, *P* < 0.001, η^2^p = 0.51). It indicated that with more options, participants fixated on a smaller fraction of the items and made fewer revisits to them. In contrast, dwell bias did not differ by set size (*F*_3,48_ = 0.26, *P* = 0.79, η^2^p = 0.016). There was a strong Demand × Set Size interaction for the screening ratio (*F*_3,48_ = 20.88, *P* < 0.001, η^2^p = 0.57): as the sets grew larger, participants inspected an even smaller proportion of options under the low-demand compared to the high-demand condition. Dwell bias also exhibited Demand × Set Size interaction: it became more negative in the low-demand condition as set size increased (*F*_3,48_ = 4.85, *P* < 0.01, η^2^p = 0.23). Since a more negative dwell bias indicates a more uneven allocation of gaze on chosen options, this pattern suggests that in easy-choice scenarios, participants zeroed in on a few promising options. There was no significant Demand × Set Size interaction for the revisit ratio (*F*_3,48_ = 2.55, *P* = 0.066, η^2^p = 0.14) (Fig. 1b, right). Notably, aggregated gaze heatmaps revealed no systematic spatial bias (i.e., participants did not consistently favor any particular screen region), confirming that differences in search behavior were due to experimental manipulations rather than layout (Supplementary Fig. 3). A similar even distribution of gaze was observed in Experiment 2 (Supplementary Fig. 4).

Neural measures also revealed a pattern of reduced effort per item in the low-demand condition. The frontal N2 latency peaked significantly earlier in the low-demand than in the high-demand conditions (*F*_1,16_ = 6.67, *P* < 0.05, η^2^p = 0.29), suggesting that decision-related conflict was resolved more rapidly for each item when the options were easier (Fig. 1c, left)^49^. Similarly, the parietal N1 and occipital P2 latencies were significantly shorter in the low-demand condition (*F*_1,16_ = 6.66, *P* < 0.05, η^2^p = 0.29; *F*_1,16_ = 5.17, *P* < 0.05, η^2^p = 0.24, respectively), indicating faster attentional orienting and perceptual-evaluative processing for each option when all options were highly attractive. Importantly, these demand-related differences varied with set size. As set size increased, all three FRP component latencies increased significantly (frontal N2: *F*_3,48_ = 23.06, *P* < 0.001, η^2^p = 0.59; parietal N1: *F*_3,48_ = 13.45, *P* < 0.001, η^2^p = 0.46; occipital P2: *F*_3,48_ = 63.26, *P* < 0.001, η^2^p = 0.80) (Fig. 1c, right). Significant Demand × Set Size interactions were observed for all three FRP latencies (frontal N2: *F*_3,48_ = 3.42, *P* < 0.05, η^2^p = 0.18; parietal N1: *F*_3,48_ = 2.94, *P* < 0.05, η^2^p = 0.16; occipital P2: *F*_3,48_ = 3.80, *P* < 0.05, η^2^p = 0.19), indicating that the divergence between low-and high-demand conditions became more pronounced as the number of options increased.

Likewise, each item fixation induced weaker occipital alpha suppression in the low-demand condition (*F*_1,16_ = 4.91, *P* < 0.05, η^2^p = 0.23) (Fig. 1d, left), indicating that participants allocated less attentional effort to each option when all choices were appealing^44–48^. Occipital alpha suppression also increased with set size (*F*_3,48_ = 13.70, *P* < 0.001, η^2^p = 0.46), reflecting higher attentional load as the assortment grew (Fig. 1d, right). In contrast, the Demand × Set Size interaction was not significant (*F*_3,48_ = 1.68, *P* = 0.20, η^2^p = 0.10), indicating that the demand-related reduction in attentional engagement was similar across set sizes. In sum, although increasing set size imposed growing neural costs per item, participants in the low-demand condition consistently processed each option with reduced cognitive effort, which became increasingly pronounced as the number of options grew.

After each choice, participants provided subjective ratings of their experience (Table 1). Paradoxically, even though the low-demand condition was rated as more enjoyable and more satisfying than the high-demand condition (*F*_1,16_ = 33.25, *P* < 0.001, η^2^p = 0.68; *F*_1,16_ = 40.20, *P* < 0.001, η^2^p = 0.72, respectively), participants reported lower confidence and higher regret in the low-demand condition (*F*_1,16_ = 22.01, *P* < 0.001, η^2^p = 0.58; *F*_1,16_ = 42.68, *P* < 0.001, η^2^p = 0.73, respectively). Set size had negligible effects on these self-reported measures (no consistent overload trend). Given the well-known limitations of post-choice self-reports, these subjective outcomes should be interpreted with caution^50–52^.

**Table 1.**
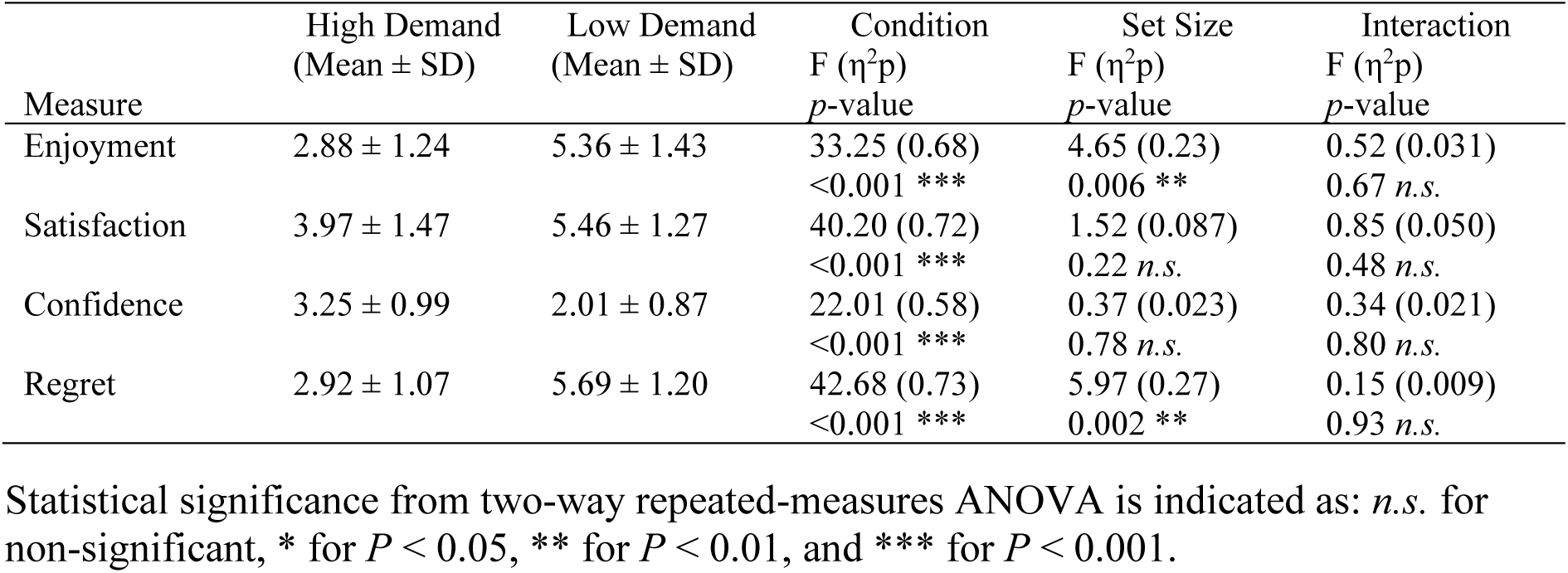
Self-reported decision experience ratings in Experiment 1.

### Experiment 2 (Sequential Narrowing)

In Experiment 2, using a bookmark-based decision aid to pre-filter options made choices significantly faster on average (*F*_1,27_ = 40.24, *P* < 0.001, η^2^p = 0.60), but did not improve choice performance. Overall, there were robust main effects of set size on choice performance (*F*_3,81_ = 33.10, *P* < 0.001, η^2^p = 0.55) and decision time (*F*_3,81_ = 63.02, *P* < 0.001, η^2^p = 0.70), reflecting a general decline in performance and increase in deliberation time as the number of options increased across both demand conditions. Yet, there was no main effect of demand on the rating difference (*F*_1,27_ = 0.21, *P* = 0.65, η^2^p = 0.010), nor any Demand × Set Size interaction (*F*_3,81_ = 0.060, *P* = 0.98, η^2^p = 0.002) (Fig. 2a, left). Unlike in Experiment 1, the Demand × Set Size interaction for decision time was not significant (*F*_3,81_ = 1.90, *P* = 0.14, η^2^p = 0.066), indicating that the aid sped up decisions by a similar amount regardless of set size (Fig. 2a, right). Thus, even with the aid, participants still took longer to choose when more options were on the screen, scaling their deliberation time with the number of items.

**Figure 2.**
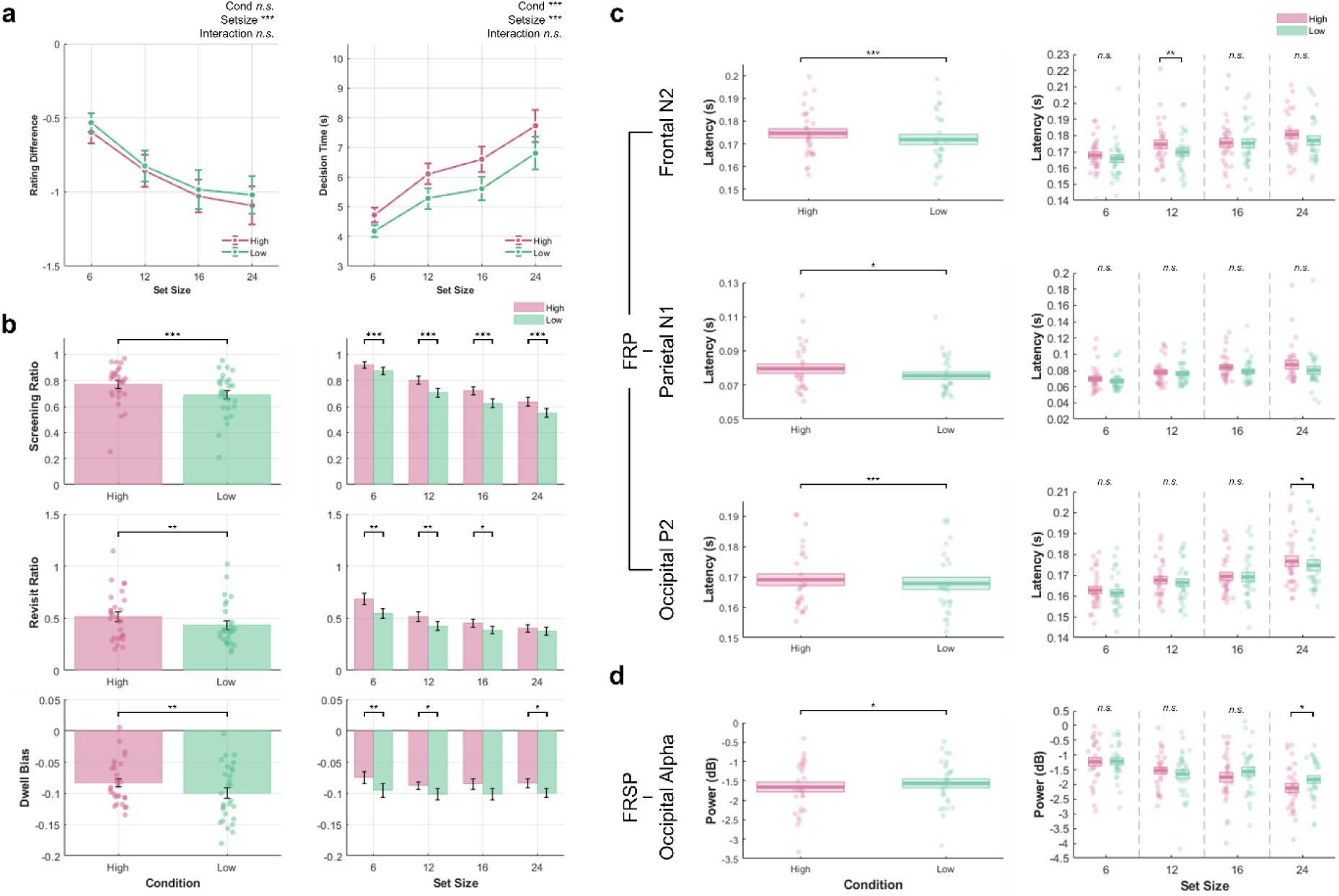
Results of Experiment 2 (Sequential narrowing). **a**, Rating difference between the chosen item and the best-rated item in the set (left) and decision time (right), shown by demand condition (high vs. low) and set size (6, 12, 16, and 24). **b**, Eye-tracking measures – screening ratio (fraction of items viewed), revisit ratio (proportion of items fixated more than once), and dwell bias (normalized gaze-time bias) – shown by demand condition (left; individual participants, n = 28) and by set size (right). **c**, EEG FRP peak latencies **d**, EEG FRSP in the alpha band. In panels c-d, individual participants are shown (n = 28), and box plots summarize the mean (center line) with boundaries indicating ±SEM. Statistical significance from two-way repeated-measures ANOVA is indicated as: *n.s.* for non-significant, * for *P* < 0.05, ** for *P* < 0.01, and *** for *P* < 0.001.

The eye-gaze measures in Experiment 2 mirrored those in Experiment 1. With the aid, participants fixated a smaller proportion of the options and made fewer back-and-forth comparisons than without the aid (Screening ratio: *F*_1,27_ = 62.13, *P* < 0.001, η^2^p = 0.70; Revisit ratio: *F*_1,27_ = 13.25, *P* = 0.001, η^2^p = 0.33; Dwell bias: *F*_1,27_ = 11.73, *P* = 0.002, η^2^p = 0.30) (Fig. 2b, left). Increasing the number of options again reduced the screening ratio and revisit ratio (*F*_3,81_ = 200.34, *P* < 0.001, η^2^p = 0.88; *F*_3,81_ = 73.65, *P* < 0.001, η^2^p = 0.73, respectively), while dwell bias showed no change with set size (*F*_3,81_ = 1.17, *P* = 0.33, η^2^p = 0.041). These effects were further qualified by significant Demand × Set Size interactions for the screening ratio (*F*_3,81_ = 7.60, *P* < 0.001, η^2^p = 0.22) and revisit ratio (*F*_3,81_ = 4.07, *P* < 0.05, η^2^p = 0.13), indicating that as set size increased, the reduction in option coverage and comparative revisits under the aided condition became more pronounced. In contrast, the Demand × Set Size interaction was not significant for dwell bias (*F*_3,81_ = 0.25, *P* = 0.80, η^2^p = 0.010). Together, these results indicate that the bookmarking step streamlined the decision process: with a self-curated shortlist, participants scanned options more efficiently and engaged in less back-and-forth deliberation (Fig. 2b, right).

The EEG results revealed parallel effects of demand reduction on processing effort. As set size increased, processing of each option slowed overall: the frontal N2 latency became significantly longer with more options (*F*_3,81_ = 15.88, *P* < 0.001, η^2^p = 0.37), as did the parietal N1 and occipital P2 latencies (*F*_3,81_ = 7.29, *P* < 0.001, η^2^p = 0.21; *F*_3,81_ = 65.02, *P* < 0.001, η^2^p = 0.71, respectively). Demand condition also influenced these neural metrics. The frontal N2 peaked significantly earlier with the aid (*F*_1,27_ = 13.72, *P* < 0.001, η^2^p = 0.34), indicating that decision-related neural processing for each item was completed more quickly with the aid (Fig. 2c, left). The parietal N1 and occipital P2 latencies were likewise shorter with the aid (*F*_1,27_ = 5.63, *P* = 0.025, η^2^p = 0.17; *F*_1,27_ = 14.44, *P* < .001, η^2^p = 0.35, respectively). In addition, each item fixation induced significantly less occipital alpha suppression when choices were pre-filtered (*F*_1,27_ = 4.51, *P* < 0.05, η^2^p = 0.14), reflecting reduced attentional load per option in the aided condition (Fig. 2d, left). Unlike in Experiment 1, however, Demand × Set Size interactions were not significant for any of the neural measures (all *P*s > 0.05), indicating that the effects of demand reduction on item-level processing speed and attentional engagement were similar across set sizes. Thus, although increasing set size similarly imposed growing neural costs per item, participants consistently responded to the decision aid by reducing effort per option across all set sizes, rather than reallocating their freed cognitive capacity to more detailed processing.

Subjective ratings in Experiment 2 mirrored those from Experiment 1 (Table 2). The low-demand condition was rated more enjoyable and more satisfying than the high-demand condition (*F*_1,27_ = 48.39, *P* < 0.001, η^2^p = 0.64; *F*_1,27_ = 38.91, *P* < 0.001, η^2^p = 0.59, respectively), but participants also reported lower confidence and higher regret (*F*_1,27_ = 21.48, *P* < 0.001, η^2^p = 0.44; *F*_1,27_ = 47.77, *P* < 0.001, η^2^p = 0.64, respectively). As in Experiment 1, set-size had minimal influence on these self-reports.

**Table 2.** Self-reported decision experience ratings in Experiment 2.

| Measure | High Demand<br>(Mean ± SD) | Low Demand<br>(Mean ± SD) | Condition | Set Size | Interaction |
| --- | --- | --- | --- | --- | --- |
| | | | F ($\eta^2p$)<br>*p*-value | F ($\eta^2p$)<br>*p*-value | F ($\eta^2p$)<br>*p*-value |
| Enjoyment | 3.75 ± 1.25 | 4.48 ± 1.20 | 48.39 (0.64)<br><0.001 *** | 11.13 (0.29)<br><0.001 *** | 1.04 (0.037)<br>0.38 *n.s.* |
| Satisfaction | 4.32 ± 1.04 | 4.84 ± 1.02 | 38.91 (0.59)<br><0.001 *** | 2.23 (0.076)<br>0.091 *n.s.* | 0.55 (0.020)<br>0.65 *n.s.* |
| Confidence | 2.43 ± 0.92 | 2.00 ± 0.66 | 21.48 (0.44)<br><0.001 *** | 3.79 (0.12)<br>0.013 * | 1.47 (0.051)<br>0.23 *n.s.* |
| Regret | 4.44 ± 0.89 | 5.17 ± 0.78 | 47.77 (0.64)<br><0.001 *** | 24.74 (0.48)<br><0.001 *** | 0.36 (0.013)<br>0.78 *n.s.* |

Statistical significance from two-way repeated-measures ANOVA is indicated as: *n.s.* for non-significant, \* for *P* < 0.05, \*\* for *P* < 0.01, and \*\*\* for *P* < 0.001.

Our results provide evidence that reducing cognitive demand – either by improving the attractiveness of options or by offering a decision aid – can change how people allocate mental effort, but not always in the intended way. Across two experiments, lowering the external cognitive demand led participants to use less internal effort, not more. Thus, when the decision-making task with abundant options was made easier, people tended to behave as cognitive misers^53^: they conserved cognitive effort rather than fully exploiting the easier task to maximize their outcomes. As a result, choice overload effects on objective performance (e.g., failing to choose the top-rated option in large sets) were still observed in the low-demand conditions, even though participants subjectively felt less overloaded.

In Experiment 1, presenting a set of all highly-attractive options did not prevent the decline in choice performance as set size increased. Instead, participants showed a strong overload pattern in the low-demand condition, and neurophysiological data indicated why: they simply did not invest as much effort per item. Similarly, in Experiment 2, using a bookmark-based shortlist made the choice objectively easier, yet participants often completed the final decision with minimal effort, expending even less effort per option than they did without the aid. Consequently, they sometimes missed the optimal choice among those favorites. Thus, lowering external demands did not automatically translate into better decision performance, because people adjusted their effort downward and still ended up with suboptimal results.

### Choice Overload Persists if Effort is Under-allocated

This pattern – where reducing external cognitive demand led participants to disengage and conserve effort, while choice performance nonetheless continued to deteriorate as set size increased – supports a nuanced view of choice overload. Even though our interventions removed much of the *objective* difficulty, overload effects in performance (choosing inferior options as set size grows) were not eliminated. This occurred not because participants did not fully engage their cognitive resources. The eye-gaze data vividly illustrated this under-engagement: as more options were available, participants’ visual scanning became more fragmented and cursory. They spent less time on each item and skipped more items entirely as set size increased, indicating that it was hard to systematically evaluate a large set – presumably because they either could not attend to everything or assumed diminishing returns from doing so.

Additionally, our EEG results showed that each item in a larger set evoked stronger effort-related brain responses on average – implying each item was more mentally taxing to process in the context of a big choice set, likely because there were more comparisons in mind and more potential conflict to resolve. Ultimately, no matter how user-friendly one makes the choice environment, having to pick one option out of many imposes a cognitive load that can degrade performance. The bottleneck can be either an inability to evaluate everything (a capacity overload) or an unwillingness to do so (strategic under-effort), but both lead to the same outcome: not choosing the global optimum.

In our experiments, it appears the dominant factor was the latter (“won’t”). Participants likely *could have* scrutinized more options or deliberated longer, but they did not because of effort–reward considerations. Our findings align with the idea that people adapt by conserving effort unless they perceive a clear benefit to expending more^21,23,54,55^.

### Effort Conservation Versus Maximization

Our results emphasize that people do not automatically use all available cognitive resources to maximize decision accuracy. Consistent with prior work, individuals often avoid cognitively demanding processing even when doing so carries a performance cost^23^. Our findings add ecological validity to that notion – even in a rich, realistic decision like picking a movie for entertainment, people exhibit this effort aversion. This behavior is not mere laziness, but a reasonable adaptation from a cost–benefit standpoint. When the reward difference between options is small, additional cognitive effort yields little payoff, reducing willingness to engage^56^. This dynamic was evident in Experiment 1. In the low-demand condition, where all options were highly attractive, and the gap between the best and an average choice was small, participants exerted minimal effort. Only when options were less appealing (and thus the “gem” carried a bigger payoff) did they ramp up effort.

Our observations coincide with prior research that decision-makers shift toward satisficing strategies when the task is easy, a pattern we observed not only with high-quality options (as in Huang & Xu, 2025) but also with the use of a filtering tool, indicating this effort conservation occurs across both content- and process-based simplifications^33^. These findings have important implications for consumer decision-making, recommender systems, and other applied choice environments: simply providing decision support or simplifying options would not automatically lead to optimal outcomes, because the human decision-maker may respond by investing even less effort.

### Insights from Eye Gaze and Brain Activity

A unique aspect of our study was the multimodal measurements of eye-tracking and EEG to examine the decision process under choice overload. These measures provided converging evidence for our interpretations and added nuance to the findings. The eye-gaze data, for instance, provided insight into how participants managed their attention when faced with many options. We observed that as choice difficulty decreased, attention became more selective rather than expansive – a pattern consistent with heuristic decision strategies. This kind of gaze behavior (skimming and re-checking only a limited set of favorites) is reminiscent of the “elimination by aspects” or other heuristic strategies described by Payne *et al.* (1988), where decision-makers quickly eliminate many options based on key aspects and then compare just a small subset in depth^57^. On the other hand, participants’ broader search in the high-demand conditions suggests they used a more compensatory strategy. Thus, the eye-tracking results confirmed a strategic shift depending on external demand: from a broad search under pressure to a narrow focus when possible.

The EEG findings reinforced our interpretation at a neural level. In particular, the frontal N2 component served as a gauge of cognitive control engagement. We saw that the frontal N2 tended to peak later whenever participants were in a situation demanding careful analysis (large sets or difficult choices), consistent with N2’s role in conflict monitoring and cognitive control^43^. Likewise, occipital alpha-band activity provided insight into visual attention and working memory. In our data, alpha desynchronization became stronger as more items were in the set, reflecting how visual working memory was taxed by keeping track of multiple items. In the low-demand conditions, weaker alpha suppression hints that participants were holding less information in mind at once – perhaps considering fewer items concurrently or processing them less deeply.

Across both experiments, subjective ratings revealed a consistent paradox. Low-demand conditions were perceived as more enjoyable and satisfying than high-demand conditions, yet they also elicited lower confidence and greater regret. This pattern supports prior findings that simplified decisions can undermine confidence, especially when the process feels too effortless^34,58^. Participants may have second-guessed their choice due to reduced deliberation, whereas more difficult tasks, though less pleasant, fostered a sense of due diligence^58^. Notably, these effects did not vary by set size, suggesting that perceived task difficulty and option quality, rather than assortment size, shaped reported confidence and satisfaction.

Our study thus contributes to the neuroscience of decision-making by linking these neural markers to the choice overload paradigm. Previous neuroimaging work (e.g., Reutskaja et al., 2018) showed that BOLD activation patterns change with choice set size; we extend those findings by showing how time-resolved brain activity at the level of individual fixations reflects the ongoing management of information^19^. This approach enables a more fine-grained examination of decision processes over time, for example, by allowing inferences about when individuals disengage from further search or commit to a choice.

### Limitations and Future Directions

Nonetheless, an important limitation is that two experiments were conducted with different participant samples, so direct comparisons between them should be made with caution. We designed each experiment to test a specific intervention separately, not to directly compare the two demand-reduction strategies head-to-head. There are also several other limitations that suggest avenues for future research. First, our sample consisted mostly of young adults in a laboratory setting. Their behavior might differ from other demographics or real-world decision contexts. For instance, older adults might have different effort strategies or experience cognitive load differently, given age-related changes in working memory^59,60^. Repeating similar experiments with more diverse age groups and in different decision domains (e.g., choosing insurance plans or retirement investments, which are notoriously complex and high-stakes) would test the generality of our findings. Next, we did not explicitly manipulate or measure participants’ motivation to be accurate. We essentially assumed that everyone was trying their best within their comfort level, but it is possible that some participants simply did not care strongly about picking the top-rated movie (especially knowing it was just for a short trailer viewing).

In conclusion, our study sheds light on the cognitive miser aspect of human decision-making in the context of choice overload. We demonstrate that reducing a task’s cognitive demand does not automatically translate into better objective decisions, because people often do not invest the extra effort they have been freed. Participants gravitated toward the path of least mental resistance, even when additional effort could have paid off. This highlights a crucial consideration for both theorists and practitioners: improving decision environments is only half the battle; the other half is ensuring that decision-makers *engage* with those improvements.

Our findings have implications for the design of recommender systems and other decision-support interfaces. Many online platforms use personalized filtering to mitigate choice overload, but our results suggest that simplification may also reduce users’ engagement with the decision. Accordingly, effective system design may require not only reducing the size of the choice set, but also sustaining deliberation among the remaining options. Interface features that support this goal include tools for rapid side-by-side comparison of top candidates and lightweight prompts when a choice is made unusually quickly, which may signal insufficient evaluation of available options. This effort-conservation tendency also has consequences in high-stakes domains such as healthcare, finance, and public policy, where decision aids are increasingly used to simplify complex choices. Although these interventions can reduce overwhelm, they may also encourage minimal processing, so decision aids and policies may need safeguards that preserve engagement (e.g., highlighting meaningful within-set differences, adding brief accountability checks, or making key trade-offs salient). Overall, simplifying choices is most likely to improve outcomes when paired with mechanisms that maintain reflective evaluation.

## Methods

### Participants

Eighteen subjects (13 female; mean age 21.61±2.81 years; age range: 18∼27) participated in Experiment 1, and a separate group of thirty subjects (16 female; mean age 23.20±3.97 years; age range: 19∼32) participated in Experiment 2. After exclusions due to eye-data corruption, the data of seventeen and twenty-eight participants were analyzed in Experiments 1 and 2, respectively. All participants had normal or corrected-to-normal vision and reported no history of neurological, psychiatric, or significant medical issues. They all provided written informed consent prior to the experiment according to the approval obtained from the Institutional Review Board of the Ulsan National Institute of Science and Technology (UNISTIRB-21-35-C) and were paid for their participation after the experiment.

### Stimuli and Materials

The choice options in both experiments were drawn from a large pool of movie posters. We compiled a database of 200 movie options, each represented by its poster image (700×1000 px). The positions of the option images on screen were arranged within a fixed 3×8 grid and randomized on each trial. For trials with fewer than 24 options, only a subset of grid positions was populated, following predefined spatial patterns that distributed items evenly across the grid while leaving the remaining positions empty. This ensured that smaller set sizes occupied the full spatial extent of the display rather than being clustered in a single region. All grids were centered on the display, and spacing between items was kept uniform. This layout strategy ensured no particular screen region was consistently advantaged or disadvantaged. Each movie poster image subtended ∼5.5°×9.7° of visual angle, with ∼2.5° gaps between images. The screen background was uniform gray. Participants were instructed that on each trial, they should freely view the options and select the movie they would most like to watch (Supplementary Fig. 1).

### Apparatus and Software

Experimental sessions took place in a quiet, dark, electromagnetically shielded booth. Participants sat ∼60 cm from a 23” LCD monitor (1920×1080 resolution, 60 Hz refresh). The eye-tracker (Tobii Pro TX300, Tobii AB, Sweden) was embedded below the monitor to record gaze at 300 Hz. A standard 9-point calibration was performed at the start and repeated as needed until the validation error was <0.5°. Head position was stabilized with an adjustable chin-and-forehead rest to maintain calibration accuracy. EEG was recorded concurrently using 31-channel wet electrodes (actiCHamp, Brain Products GmbH, Germany). Electrodes were placed according to the international 10–20 system, with additional ground (left mastoid) and reference (right mastoid) electrodes. Impedances were kept below 10 kΩ throughout the recording, and the signals were sampled at 500 Hz. The stimulus presentation and data synchronization were controlled by custom scripts in MATLAB (MathWorks, Natick, MA, USA). Precise timing of stimulus presentation updates and event markers was synchronized with EEG signals using the parallel port trigger output to the EEG system.

### Experimental Design

We conducted two separate but conceptually parallel experiments, each employing a 2 × 4 within-subjects factorial design. The factors were Demand (high-demand vs. low-demand) and Set Size (number of options: 6, 12, 16, or 24). While 18 alternatives could be a plausible intermediate level between 12 and 24, we used a set size of 16 to match the consistency with the stimulus-generation method and counterbalancing constraints. Specifically, each demand condition drew from a fixed pool of 48 items – the least common multiple of the set sizes – allowing the stimulus-generation algorithm to partition the pool into balanced subsets for each set size and to populate a fixed 3×8 grid using predefined position templates with even spatial sampling. In both experiments, each participant completed choice trials under both demand conditions and all set sizes, allowing repeated-measures comparisons. Experiment 1 manipulated cognitive demand through the choice set content, whereas Experiment 2 manipulated demand through the decision process.

In Experiment 1, the low-demand condition presented choices from high-attractiveness sets (all options with high initial appeal), while the high-demand condition used low-attractiveness sets (options of lower appeal). The high-attractiveness pool comprised the 48 highest-rated posters based on each participant’s initial ratings (see below for the rating procedure), and we computed the mean and variance of ratings within this pool. The low-attractiveness pool was then selected from the remaining posters using an optimization procedure that searched candidate 48-item subsets and chose the subset that maximized the mean rating difference from the high-attractiveness pool while keeping the rating dispersion comparable. This ensured the two pools had similar rating variance while maximizing the overall rating difference.

In Experiment 2, participants were provided a bookmarking tool (decision aid) to narrow the options before the choice task. In the low-demand condition, the final choice set was constructed from a screened pool anchored on the participant’s bookmarked items, whereas in the high-demand condition, the final set was drawn from a control pool containing only non-bookmarked items. Because the number of bookmarks varied across participants, we utilized an algorithmic procedure to ensure that the screened and control pools were both fixed to 48 items and matched in rating distribution. Specifically, if a participant bookmarked more than 48 items, the screened pool was defined as the 48 highest-rated bookmarked items. If fewer than 48 bookmarked items were available, the screened pool was completed by adding non-bookmarked items selected from the highest available rating values until the pool reached 48 items (i.e., fillers were drawn preferentially from the top of the remaining rating-ranked non-bookmarked candidates; selection within a given rating value was randomized). On average, 18.93 bookmarked items (SD = 8.06) were included in the 48-item screened pool. To ensure that any differences between conditions reflected bookmarking rather than option quality, we matched the rating distribution of the screened and control pools. Specifically, after the screened pool was finalized, we then constructed the control pool by sampling exclusively from non-bookmarked items such that, for each rating bin, the control pool contained the same number of items as the screened pool. When multiple candidates were available within a bin, items were selected at random. Thus, the screened and control pools were identical in rating distribution and differed only in whether items had been bookmarked. Given the smallest set size of 6, we required that the screened pool contain at least 16.67% bookmarked items to ensure that each screened display could include at least one bookmarked option. Because 16.67% of 48 corresponds to 8 items, participants would have been excluded if fewer than 8 bookmarked items were available for the screened pool; no participants were excluded based on this criterion.

Within each experiment, trial conditions were counterbalanced and randomized. In both Experiment 1 and 2, all trials were intermixed in random order (since no additional instruction difference was needed between conditions). Each participant experienced every combination of Demand and Set Size an equal number of times. Specifically, each of the 8 conditions (2 demand × 4 set sizes) was presented 20 times, yielding 160 choice trials per participant in each experiment. Additionally, Forced Choice (FC) trials were included as attention-check control trials in which participants were instructed to choose the highlighted item regardless of preference, allowing us to verify attention and task compliance under identical visual displays. FC trials were presented 4 times for each of the 8 conditions (2 demand × 4 set sizes), yielding 32 FC trials and a total of 192 trials for all conditions per participant. The trials were separated into 4 blocks considering individuals’ fatigue. Each block consisted of the same number of trials for each condition, which would result in 48 trials (2 demand × 4 set sizes × 5 trials + 1 FC × 8 trials). Trial order within blocks was randomized for each participant. Within each trial, no item was shown more than once, and stimulus scheduling was used to distribute repeated appearances of posters approximately evenly across trials within each condition and set size, minimizing systematic familiarity differences between conditions.

### Tasks

Each participant completed a single session (∼3 hours including setup). The sequence of tasks was as follows:

- **Task 1: Familiarity Check.** Participants were shown 200 movie posters one by one and asked if they had watched the movie before. Responses were obtained in binary options (1 = I have watched the movie before, 0 = I have never watched the movie before). Only the ones they had not watched before were used for further tasks. On average, participants contributed 189.00 (SD = 10.64) eligible posters in Experiment 1 and 185.21 (SD = 11.55) eligible posters in Experiment 2 after this familiarity screening.
- **Task 2: Rating Task.** Participants were shown the unwatched movies one by one again. This time, participants rated how much they wanted to watch the movie by clicking the rating bar below the poster image. An 11-point Likert scale was used for the rating bar (1 = I do not want to watch the movie at all, 11 = I want to watch the movie very much). All stimuli were presented in random order. After completing one round of the rating task, participants went through another identical round, also in random order, making a total of two rounds of the rating task. The mean rating of each stimulus was calculated by averaging the two ratings.
- **Task 2-1: Bookmark Task.** This task was done only for Experiment 2. Participants went through a bookmark task right after the rating task. A maximum of 24 posters were shown on the screen while participants freely clicked on those they would like to bookmark. A heart-shaped icon appeared above the selected poster, indicating a bookmark. Multiple selection was allowed, and the task was self-paced until all unseen posters from Task 1 were presented.
- **Task 3: Choice Task.** The objective of the choice task was to choose the most desirable option. Participants were informed that they would watch a short review of one of their chosen movies after the experiment. Each participant completed 4 blocks of 48 trials each with a short break between the blocks. Choices were self-paced with no explicit time limit; the trial ended when a selection was made.
- **Task 4: Post-Choice Ratings.** After each choice trial in both experiments, participants answered a brief set of self-report questions about their decision experience (while the EEG/eye-tracking recording continued passively during this period). Specifically, they rated: (a) how enjoyable the decision process was, (b) how satisfied they were with their chosen option, (c) how confident they were that they made the best choice, and (d) how much regret they felt. Ratings were made on 7-point Likert scales presented on the screen (tailored to each question, e.g., 1 = not at all enjoyable, 7 = extremely enjoyable). Participants pressed number keys to submit their responses. Once the questions were completed, the next choice trial began.

Throughout the procedure, we monitored EEG and eye-tracking signals in real-time for quality. If drift or signal loss was detected (e.g., due to cap shift or calibration error), the experiment was paused to recalibrate the eye-tracker or adjust electrodes. Such pauses were infrequent, and data collection resumed once signals were stable.

### Task Structure

Each choice trial followed a fixed timeline. A central fixation cross appeared at trial start for 2 s, giving participants a moment to focus and blank out previous visual stimuli. The choice set then appeared, and participants began exploring the options freely. In all conditions, the choice set display remained on-screen until a decision was made (self-paced). Participants indicated their choice by clicking their chosen item using a mouse. The time from set onset to mouse click constituted the decision time measure for that trial. Once a choice was made, the screen cleared, and the four experience-rating questions (Task 4) were presented one by one at the center. Participants responded to each within a few seconds (self-paced), and this pacing ensured participants had a short break to relax their eyes between trials. The full experimental procedure and specific manipulations for each demand condition are depicted in Supplementary Fig. 1.

### Eye-Tracking Acquisition and Preprocessing

Eye-tracking data were processed to extract gaze fixation events and visual attention measures. A fixation was defined as a period during which gaze remained within an Area of Interest (AOI) for at least 100 ms. Brief data gaps (<100 ms) were bridged through linear interpolation. After detecting fixations, we mapped each fixation to a specific choice option or to “blank” (if the fixation fell on background areas). Only fixations on actual option items were retained for effort analyses. Each option’s on-screen image was defined as an AOI with a bounding box slightly larger than the image to capture near-boundary gaze points.

From the sequence of fixations on each trial, we computed several eye gaze metrics characterizing visual search behavior^36–39^. First, the Screening Ratio was calculated as the proportion of available options that the participant fixated on at least once, given by:

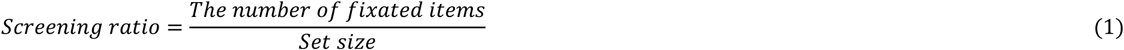

This measures how exhaustive the search was: a screening ratio of 1 means the person looked at every option, whereas lower values indicate some items were never seen. Second, the Revisit Ratio indexed comparative look-backs. For each trial, we computed the average number of times the participant returned to an option after initially seeing it, given by:

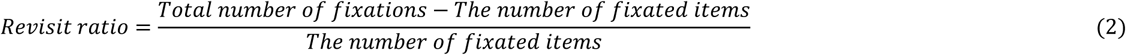

A higher revisit ratio means that options were viewed multiple times on average (suggesting more back-and-forth comparing), whereas a value of 0 would mean a pure single-pass scan with no returns. Third, we quantified gaze concentration using a Dwell Time Inequality measure, termed Dwell Bias. For each trial, we compared the proportion of total decision time spent fixating on unchosen options with the proportion of fixated options that were unchosen as follows:

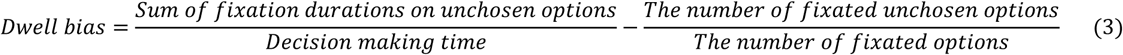

All options except the eventually chosen one were considered unchosen options. Dwell bias can take positive or negative values and is centered at 0. It becomes 0 when the proportion of total fixation time devoted to unchosen options equals the proportion of uniquely fixated options that were unchosen – that is, when gaze time is allocated to unchosen items in direct proportion to how many unchosen items were inspected. Negative dwell bias indicates that unchosen options received less fixation time than expected, given how many were viewed, consistent with attention being concentrated on the chosen option rather than distributed across the unchosen options. We used this measure to capture whether participants focused disproportionately on their top candidate (which can reflect a narrow decision strategy). Together, screening ratio, revisit ratio, and dwell bias provide a comprehensive profile of visual search strategy – from breadth of exploration to depth of comparative examination.

### EEG Acquisition and Preprocessing

EEG data were preprocessed and analyzed using the EEGLAB toolbox and custom MATLAB scripts. Continuous EEG recordings were first high-pass filtered (0.1 Hz) to remove slow drifts. A notch filter at 60 Hz was applied to attenuate line noise. Then, a band-pass filter from 1 to 50 Hz was applied (finite impulse response filter, zero-phase). Bad channels were detected using kurtosis-based criteria and were interpolated using spherical spline interpolation (on average 1–2 channels per participant)^61,62^. After that, the signals were re-referenced by the common average reference (CAR) method. We performed an Independent Component Analysis (ICA) to identify and remove residual oculomotor artifacts. The data were then processed with an Artifact Subspace Reconstruction (ASR) method to remove high-amplitude transient artifacts. We used a conservative threshold of 20 standard deviations for the ASR cutoff, which has been shown to effectively suppress large artifacts while preserving genuine EEG signals^63^. Next, we segmented the EEG into epochs time-locked to fixation onsets.

We extracted fixation-locked EEG epochs for each option fixation during the choice tasks. Using the eye-tracking data, we took each instance when the participant fixated on an option (AOI) and marked that timestamp in the EEG. Epochs were defined from 200 ms before fixation onset to 800 ms after onset. Baseline correction was applied using a different baseline onset. Since each fixation-onset baseline could overlap with the previous fixation, all fixation epochs in a trial were corrected using a common baseline by subtracting the mean pre-trial range (−200 to 0 ms) from the trial-onset for each channel^64^. Note that a post-hoc baseline correction (correcting the baseline range again) was applied to match the value of the baseline range to zero, only for the visualization purpose. This only induced a vertical shift in absolute values for visual convenience, and all other analyses (including statistical tests) were performed without the post-hoc baseline correction. These fixation-aligned EEG epochs form the basis of our neural measures of processing effort.

For each participant and each experimental condition, we averaged the fixation-locked EEG epochs to compute Fixation-Related Potentials (FRPs). FRPs are analogous to traditional event-related potentials, but time-locked to the onset of each fixation on an option. We focused on three FRP components that have been linked to visual and cognitive processing in decision-making: an early posterior N1, a mid-latency occipital P2, and a later frontal N2. These components were identified in the grand-average waveform and by prior literature^40,41,43^. The N1 was defined as the first prominent negative deflection around 100 ms, maximal at parietal-occipital sites (we measured it at a pooled electrode cluster around Pz/P3/P4). The P2 was a subsequent positive deflection at ∼200 ms, prominent at occipital sites (measured at Oz/O1/O2). The N2 was a negative deflection ∼200 ms post-fixation, strongest at frontal sites (measured at Fz/F3/F4). For each participant in each condition, we extracted the peak latency of each of these components: specifically, the time point of the most negative amplitude of N1 within 0–200 ms (N1 latency), the most positive of P2 within 100–250 ms (P2 latency), and most negative of N2 within 100–250 ms (N2 latency). These latencies served as neural measures of processing speed for each item: longer latencies indicate longer processing of the visual stimulus, consistent with more effort.

In addition to time-domain FRPs, we analyzed the EEG in the time-frequency domain to measure changes in oscillatory activity in response to fixated stimuli. We focused on oscillatory activity in the alpha frequency band (8–13 Hz) over visual cortex, as alpha power reductions are a known index of attentional engagement^44–48^. For each fixation epoch, we computed a wavelet-based time-frequency decomposition at occipital sites (Oz, O1, O2). From alpha-band power time-course, we extracted the alpha desynchronization magnitude, defined as the largest decrease in alpha power within 0–300 ms after fixation onset relative to a common pre-trial baseline (−200 to 0 ms relative to trial onset). This reflects how much the occipital alpha-band activity was suppressed when the option was being processed. A larger alpha suppression (more negative change) indicates stronger visual cortical activation and attention to the stimulus. We denote this neural measure as FRSP (with more negative values corresponding to greater attentional effort).

In summary, our EEG analysis yielded two classes of neural metrics for each participant and condition: (1) FRP component latencies (N1, P2, N2) as indicators of processing speed per option, and (2) FRSP alpha suppression magnitude as an indicator of attentional intensity per option. These were computed for all conditions in both experiments, allowing us to assess how cognitive demand and set size affected the depth and speed of neural processing for each option encountered.

### Statistical Analysis

All statistical analyses were conducted using MATLAB with the Statistics and Machine Learning Toolbox. For each experiment, we used repeated-measures ANOVA to test the effects of Demand and Set Size on the outcome measures (i.e., dependent variables). Specifically, for each measure (i.e., choice performance, decision time, each eye gaze measure, each EEG measure), we fit a two-way within-subjects ANOVA model with Demand (low vs. high) and Set Size (6, 12, 16, 24) as factors. The ANOVAs were implemented via MATLAB’s fitrm and ranova functions, which produce a full factorial repeated-measures analysis. An alpha level of 0.05 (two-tailed) was used for all significance tests. We report F-statistics with numerator/denominator degrees of freedom and P-values. Effect sizes are given as partial eta squared (η^2^p). Post-hoc pairwise comparisons were conducted using paired t-tests with Bonferroni correction to control for multiple comparisons.

### Data and Code Availability

All data, materials, and analysis code for this study are available from the corresponding author upon reasonable request.

## Acknowledgements

This work was supported by the National Research Foundation of Korea (NRF) grant funded by the Korea government (MSIT) (RS-2025-00522357) and the IITP (Institute of Information & Communications Technology Planning & Evaluation) - ITRC (Information Technology Research Center) grant funded by the Korea government (Ministry of Science and ICT) (IITP-2026-RS-2024-00437866).

## References

1. Wansink, B. & Sobal, J. Mindless Eating: The 200 Daily Food Decisions We Overlook. Environment and Behavior 39, 106–123 (2007).

2. Broniarczyk, S. M., Hoyer, W. D. & McAlister, L. Consumers’ Perceptions of the Assortment Offered in a Grocery Category: The Impact of Item Reduction. Journal of Marketing Research 35, 166–176 (1998).

3. Kahn, B. E. & Lehmann, D. R. Modeling Choice Among Assortments.

4. Kahn, B., Moore, W. L. & Glazer, R. Experiments in Constrained Choice. Journal of Consumer Research 14, 96–113 (1987).

5. Babin, B. J., Darden, W. R. & Griffin, M. Work and/or Fun: Measuring Hedonic and Utilitarian Shopping Value. Journal of Consumer Research 20, 644–656 (1994).

6. Inbar, Y., Botti, S. & Hanko, K. Decision speed and choice regret: When haste feels like waste. Journal of Experimental Social Psychology 47, 533–540 (2011).

7. Zeelenberg, M. et al. Emotional Reactions to the Outcomes of Decisions: The Role of Counterfactual Thought in the Experience of Regret and Disappointment. Organizational Behavior and Human Decision Processes 75, 117–141 (1998).

8. Botti, S. & lyengar, S. S. The Psychological Pleasure and Pain of Choosing: When People Prefer Choosing at the Cost of Subsequent Outcome Satisfaction. Journal of Personality and Social Psychology 87, 312–326 (2004).

9. Dhar, R. & Nowlis, S. M. The Effect of Time Pressure on Consumer Choice Deferral. Journal of Consumer Research 25, 369–384 (1999).

10. Iyengar, S. S. & Lepper, M. R. When choice is demotivating: Can one desire too much of a good thing? Journal of Personality and Social Psychology 79, 995–1006 (2000).

11. Chernev, A., Böckenholt, U. & Goodman, J. Choice overload: A conceptual review and meta-analysis. Journal of Consumer Psychology 25, 333–358 (2015).

12. Kim, H., Choi, J. H. & Bao, T. Why Does Netflix Syndrome Occur: A Study on the Effect of Content Choice Deferral on Stress. AJPOR 13, (2025).

13. Hauser, J. R. & Wernerfelt, B. An Evaluation Cost Model of Consideration Sets. Journal of Consumer Research 16, 393–408 (1990).

14. Huffman, C. & Kahn, B. E. Variety for sale: Mass customization or mass confusion? Journal of Retailing 74, 491–513 (1998).

15. Jacoby, J., Speller, D. E. & Kohn, C. A. Brand Choice Behavior as a Function of Information Load. Journal of Marketing Research 11, 63–69 (1974).

16. Malhotra, N. K. Information Load and Consumer Decision Making. Journal of Consumer Research 8, 419–430 (1982).

17. Scammon, D. ‘Information Load’ and Consumers. Journal of Consumer Research 4, 148–55 (1977).

18. Shugan, S. M. The Cost of Thinking. Journal of Consumer Research 7, 99–111 (1980).

19. Reutskaja, E., Lindner, A., Nagel, R., Andersen, R. & Camerer, C. Choice overload reduces neural signatures of choice set value in dorsal striatum and anterior cingulate cortex. Nature Human Behaviour 2, (2018).

20. Mækelæ, M. J. et al. Is it cognitive effort you measure? Comparing three task paradigms to the Need for Cognition scale. PLOS ONE 18, e0290177 (2023).

21. Westbrook, A. & Braver, T. S. Cognitive effort: A neuroeconomic approach. Cogn Affect Behav Neurosci 15, 395–415 (2015).

22. Hull, C. L. Principles Of Behavior. (1943).

23. Kool, W., McGuire, J. T., Rosen, Z. B. & Botvinick, M. M. Decision making and the avoidance of cognitive demand. J Exp Psychol Gen 139, 665–682 (2010).

24. Westbrook, A., Kester, D. & Braver, T. S. What Is the Subjective Cost of Cognitive Effort? Load, Trait, and Aging Effects Revealed by Economic Preference. PLOS ONE 8, e68210 (2013).

25. Sweller, J. Cognitive load during problem solving: Effects on learning. Cognitive Science 12, 257–285 (1988).

26. Misuraca, R., Nixon, A. E., Miceli, S., Di Stefano, G. & Scaffidi Abbate, C. On the advantages and disadvantages of choice: future research directions in choice overload and its moderators. Front Psychol 15, 1290359 (2024).

27. Chernev, A. & Hamilton, R. Assortment Size and Option Attractiveness in Consumer Choice among Retailers. Journal of Marketing Research 46, 410–420 (2009).

28. Corcoran, K. & Mussweiler, T. The cognitive miser’s perspective: Social comparison as a heuristic in self-judgements. European Review of Social Psychology 21, 78–113 (2010).

29. Genevsky, A. & Yoon, C. Neural basis of consumer decision making and neuroforecasting. in APA handbook of consumer psychology. 621–635 (American Psychological Association, Washington, 2022). doi:10.1037/0000262-027.

30. Costa-Feito, A., González-Fernández, A. M., Rodríguez-Santos, C. & Cervantes-Blanco, M. Electroencephalography in consumer behaviour and marketing: a science mapping approach. Humanit Soc Sci Commun 10, 474 (2023).

31. Peng, M., Xu, Z. & Huang, H. How Does Information Overload Affect Consumers’ Online Decision Process? An Event-Related Potentials Study. Front. Neurosci. 15, (2021).

32. Hu, X., Meng, Z. & He, Q. Choice overload interferes with early processing and necessitates late compensation: Evidence from electroencephalogram. European Journal of Neuroscience 59, 2995–3008 (2024).

33. Huang, X. & Xu, S. Mitigating choice overload: The interactive effects of set size and overall preference revealed by hierarchical drift diffusion modeling and electroencephalography. NeuroImage 321, 121542 (2025).

34. Chernev, A. When More Is Less and Less Is More: The Role of Ideal Point Availability and Assortment in Consumer Choice. J Consum Res 30, 170–183 (2003).

35. Shah, A. M. & Wolford, G. Buying Behavior as a Function of Parametric Variation of Number of Choices. Psychol Sci 18, 369–370 (2007).

36. Gidlöf, K., Wallin, A., Dewhurst, R. & Holmqvist, K. Using Eye Tracking to Trace a Cognitive Process: Gaze Behaviour During Decision Making in a Natural Environment. Journal of Eye Movement Research 6, 1–14 (2013).

37. Wiegand, I., van Pouderoijen, M., Oosterman, J. M., Deckers, K. & Horstmann, G. Contributions of distractor dwelling, skipping, and revisiting to age differences in visual search. Sci Rep 15, 1801 (2025).

38. Horstmann, G., Herwig, A. & Becker, S. I. Distractor Dwelling, Skipping, and Revisiting Determine Target Absent Performance in Difficult Visual Search. Front. Psychol. 7, (2016).

39. Becker, S. I. Determinants of Dwell Time in Visual Search: Similarity or Perceptual Difficulty? PLoS One 6, e17740 (2011).

40. Haider, M., Spong, P. & Lindsley, D. B. Attention, Vigilance, and Cortical Evoked-Potentials in Humans. Science 145, 180–182 (1964).

41. Cepeda-Freyre, H. A., Garcia-Aguilar, G., Eguibar, J. R. & Cortes, C. Brain Processing of Complex Geometric Forms in a Visual Memory Task Increases P2 Amplitude. Brain Sci 10, 114 (2020).

42. Folstein, J. R. & Van Petten, C. Influence of cognitive control and mismatch on the N2 component of the ERP: A review. Psychophysiology 45, 152–170 (2008).

43. Xie, L., Cao, B., Li, Z. & Li, F. Neural Dynamics of Cognitive Control in Various Types of Incongruence. Front. Hum. Neurosci. 14, (2020).

44. Hanslmayr, S., Staudigl, T. & Fellner, M.-C. Oscillatory power decreases and long-term memory: the information via desynchronization hypothesis. Front. Hum. Neurosci. 6, (2012).

45. Hanslmayr, S., Staresina, B. P. & Bowman, H. Oscillations and Episodic Memory: Addressing the Synchronization/Desynchronization Conundrum. Trends in Neurosciences 39, 16–25 (2016).

46. Griffiths, B. J. et al. Alpha/beta power decreases track the fidelity of stimulus-specific information. eLife 8, e49562 (2019).

47. Gevins, A. et al. Monitoring Working Memory Load during Computer-Based Tasks with EEG Pattern Recognition Methods. Hum Factors 40, 79–91 (1998).

48. Fairclough, S. H. & Ewing, K. The effect of task demand and incentive on neurophysiological and cardiovascular markers of effort. International Journal of Psychophysiology 119, 58–66 (2017).

49. Lin, Y.-Q. et al. N1 and P1 Components Associate With Visuospatial-Executive and Language Functions in Normosmic Parkinson’s Disease: An Event-Related Potential Study. Front. Aging Neurosci. 11, (2019).

50. Nisbett, R. E. & Wilson, T. D. Telling more than we can know: Verbal reports on mental processes. Psychological Review 84, 231–259 (1977).

51. Schwarz, N. Self-reports: How the questions shape the answers. American Psychologist 54, 93–105 (1999).

52. Wilson, T. D. & Dunn, E. W. Self-Knowledge: Its Limits, Value, and Potential for Improvement. Annu. Rev. Psychol. 55, 493–518 (2004).

53. Fiske, S. T. & Taylor, S. E. Social Cognition. (McGraw-Hill Ryerson, Limited, 1984).

54. Shenhav, A., Botvinick, M. M. & Cohen, J. D. The Expected Value of Control: An Integrative Theory of Anterior Cingulate Cortex Function. Neuron 79, 217–240 (2013).

55. Kurzban, R., Duckworth, A., Kable, J. W. & Myers, J. An opportunity cost model of subjective effort and task performance. Behavioral and Brain Sciences 36, 661–679 (2013).

56. Apps, M. A. J., Grima, L. L., Manohar, S. & Husain, M. The role of cognitive effort in subjective reward devaluation and risky decision-making. Sci Rep 5, 16880 (2015).

57. Payne, J. W., Bettman, J. R. & Johnson, E. J. Adaptive strategy selection in decision making. *Journal of Experimental Psychology: Learning*, Memory, and Cognition 14, 534–552 (1988).

58. Kruger, J., Wirtz, D., Van Boven, L. & Altermatt, T. W. The effort heuristic. Journal of Experimental Social Psychology 40, 91–98 (2004).

59. Bopp, K. L. & Verhaeghen, P. Aging and Verbal Memory Span: A Meta-Analysis. J Gerontol B Psychol Sci Soc Sci 60, P223–P233 (2005).

60. Park, D. C. & Reuter-Lorenz, P. The Adaptive Brain: Aging and Neurocognitive Scaffolding. Annual Review of Psychology 60, 173–196 (2009).

61. Delorme, A. & Makeig, S. EEGLAB: an open source toolbox for analysis of single-trial EEG dynamics including independent component analysis. Journal of Neuroscience Methods 134, 9–21 (2004).

62. Delorme, A., Sejnowski, T. & Makeig, S. Enhanced detection of artifacts in EEG data using higher-order statistics and independent component analysis. NeuroImage 34, 1443–1449 (2007).

63. Chang, C.-Y., Hsu, S.-H., Pion-Tonachini, L. & Jung, T.-P. Evaluation of Artifact Subspace Reconstruction for Automatic EEG Artifact Removal. in 2018 40th Annual International Conference of the IEEE Engineering in Medicine and Biology Society (EMBC) 1242–1245 (2018). doi:10.1109/EMBC.2018.8512547.

64. Nikolaev, A. R., Meghanathan, R. N. & van Leeuwen, C. Combining EEG and eye movement recording in free viewing: Pitfalls and possibilities. Brain and Cognition 107, 55–83 (2016).

