## Supplementary Information for "Effects of Cognitive Demand Reduction on Choice Overload"

### S1. Supplementary Methods

#### S1.1. Experimental Paradigm

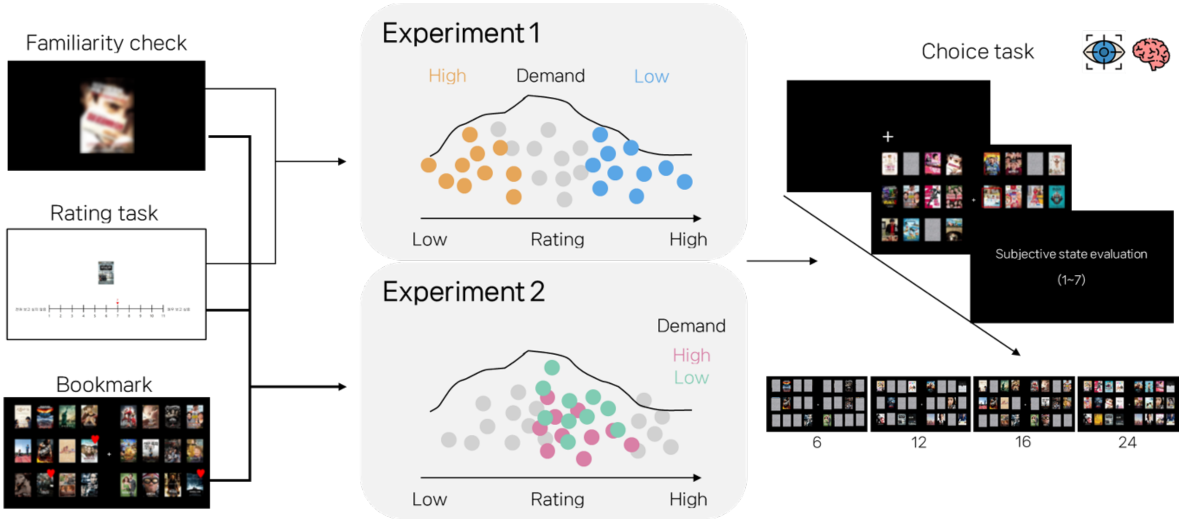

**Supplementary Fig. 1 | Experimental procedure and demand manipulations**

Schematic of task sequence and demand conditions in Experiments 1 and 2. Participants first completed a familiarity screening and then rated movie posters for attractiveness (11-point scale). **Experiment 1 (Preference-based Filtering):** High-demand condition used a pool of low-rated posters, and low-demand condition used high-rated posters (so that “easier” choices had uniformly high-attractiveness options). **Experiment 2 (Sequential Narrowing):** Participants bookmarked favorites after rating; in low-demand trials, the final choice set included bookmarked items, whereas in high-demand trials, the set was drawn from a rating-matched control pool without bookmarks. On each choice trial, participants saw an array of 6, 12, 16, or 24 posters, selected one, and then rated their decision experience (7-point scale). Eye-gaze and EEG were recorded during the choice task.

#### S1.2. Manipulation check

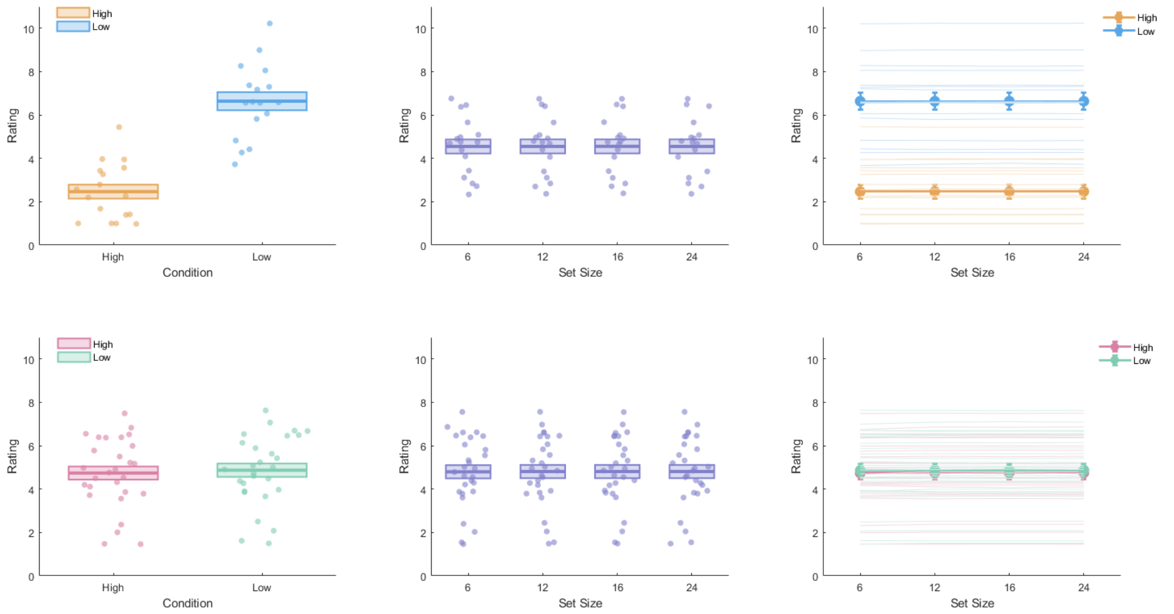

**Supplementary Fig. 2 | Manipulation check**

Plots show the mean (±SEM) rating of the stimulus pools used to construct the choice sets under each demand condition for Experiments 1 and 2 (ratings obtained from each participant's initial rating task on an 11-point scale). In Experiment 1, the high-attractiveness (low-demand) pool was rated higher than the low-attractiveness (high-demand) pool, confirming the intended manipulation (two-way repeated-measures ANOVA with Demand and Set Size: main effect of Demand, $F_{1,16} = 1129.6$, $P < 0.001$, $\eta^2 p = 0.89$; main effect of Set Size, $F_{1,16} = 0.17$, $P = 0.75$, $\eta^2 p = 0.011$; Demand × Set Size, $F_{3,48} = 0.48$, $P = 0.55$, $\eta^2 p = 0.029$). In Experiment 2, the screened (low-demand) and matched control (high-demand) pools were constructed to have closely similar attractiveness; accordingly, the observed mean difference between conditions was small ($\Delta = 0.13$ on the 11-point scale). Although the ANOVA detected a statistically significant main effect of Demand ($F_{1,27} = 251.54$, $P < 0.001$, $\eta^2 p = 0.90$), the magnitude of the difference was minimal in absolute terms, consistent with the goal of equating option quality across conditions while varying whether choice sets contained bookmarked items. The main effect of Set Size was ($F_{1,27} = 2.99$, $P = 0.009$, $\eta^2 p = 0.010$), and the Demand × Set Size interaction was ($F_{3,81} = 0.37$, $P = 0.59$, $\eta^2 p = 0.014$), indicating that any residual difference between conditions did not systematically vary across set sizes.

### S2. Supplementary Results

#### S2.1. Spatial-bias check

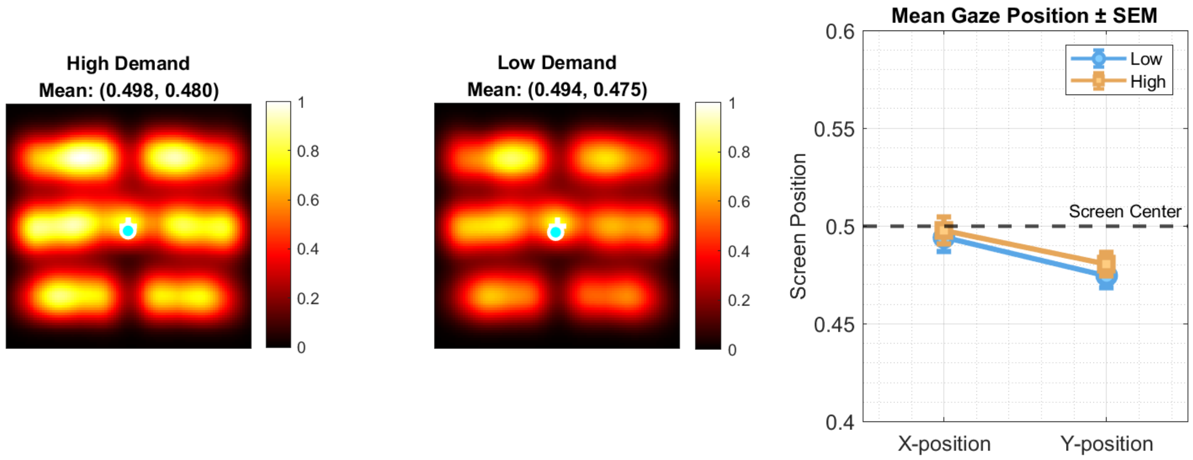

**Supplementary Fig. 3 | Eye-gaze heatmaps and mean gaze position show no systematic spatial viewing bias (Experiment 1)**

The left panel shows aggregated gaze heatmaps for the high- and low-demand conditions; the overlaid marker denotes the mean gaze position (reported as normalized screen coordinates, x and y). Both conditions showed highly similar gaze distributions, with mean (±SEM) gaze positions near the screen center (High-demand mean = (0.498, 0.480); Low-demand mean = (0.494, 0.475)). The right panel shows the mean gaze position collapsed across trials for each axis (x-position and y-position), plotted relative to the screen center (dashed line at 0.5). To quantify potential spatial bias, mean gaze position was analyzed separately for x and y coordinates using two-way repeated-measures ANOVAs. There was no evidence of systematic spatial bias for both x- and y-positions (x-position: $F_{1,16} = 2.09$, $P = 0.17$, $\eta^2p = 0.12$; y-position: $F_{1,16} = 1.25$, $P = 0.28$, $\eta^2p = 0.072$).

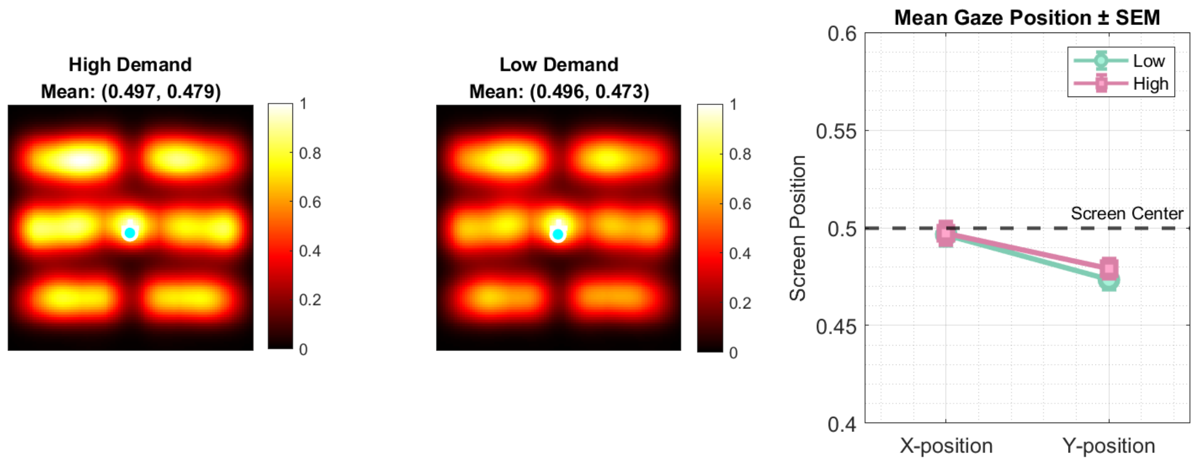

**Supplementary Fig. 4 | Eye-gaze heatmaps and mean gaze position show no systematic spatial viewing bias (Experiment 2)**

The left panel shows aggregated gaze heatmaps for the high- and low-demand conditions; the overlaid marker denotes the mean gaze position (reported as normalized screen coordinates, x and y). Both conditions showed highly similar gaze distributions, with mean gaze positions near the screen center (High-demand mean = (0.497, 0.479); Low-demand mean = (0.496, 0.473)). The right panel shows the mean gaze position collapsed across trials for each axis (x-position and y-position), plotted relative to the screen center (dashed line at 0.5). To quantify potential spatial bias, mean gaze position was analyzed separately for x and y coordinates using two-way repeated-measures ANOVAs. There was no evidence of systematic spatial bias for both x- and y-positions (x-position: $F_{1,27} = 0.036$, $P = 0.85$, $\eta^2p = 0.001$; y-position: $F_{1,27} = 1.33$, $P = 0.26$, $\eta^2p = 0.047$).
